# Full Title: Growth substrate may influence biofilm susceptibility to antibiotics

**DOI:** 10.1101/449892

**Authors:** Dustin L. Williams, Scott R. Smith, Brittany R. Peterson, Gina Allyn, Richard Tyler Epperson, Ryan E. Looper

## Abstract

The CDC biofilm reactor is a robust culture system with high reproducibility in which biofilms can be grown for a wide variety of analyses. Multiple material types are available as growth substrates, yet data from biofilms grown on biologically relevant materials is scarce, particularly for antibiotic efficacy against differentially supported biofilms. In this study, CDC reactor holders were modified to allow growth of biofilms on collagen, a biologically relevant substrate. Susceptibility to multiple antibiotics was compared between biofilms of varying species grown on collagen versus standard polycarbonate coupons. Data indicated that in 13/18 instances, biofilms on polycarbonate were more susceptible to antibiotics than those on collagen, suggesting that when grown on a complex substrate, biofilms may be more tolerant to antibiotics. These outcomes may influence the translatability of antibiotic susceptibility profiles that have been collected for biofilms on hard plastic materials. Data may also help to advance information on antibiotic susceptibility testing of biofilms grown on biologically relevant materials for future *in vitro* and *in vivo* applications.

## Introduction

The CDC biofilm reactor has been validated as a robust and repeatable reactor system for elucidating various aspects of biofilm physiology, morphology, growth dynamics and antibiotic susceptibility profiles (Buckingham-Meyer et al. 2007, Gilmore et al. 2010, Goeres et al. 2005, Honraet et al. 2005, Williams, Woodbury, et al. 2011). To analyze biofilms, their characteristics and properties, the design of the CDC reactor system is such that coupons can be placed in holding rods (Figure 1A). This allows for exposure to shear force and renewable nutrients that optimize biofilm formation by simulating natural environments such as rivers, streams, oral cavities, biomedical device surfaces or industrial systems (Costerton et al. 1978, Costerton and Irvin 1981, Darouiche 2004, Gibbons and Houte 1975, Stoodley et al. 2001). To mimic such environments, coupons are manufactured from a wide variety of materials. For example, to simulate the surface of culinary water pipelines, coupons made of iron-based metals or polyvinylchloride (PVC) are available (BiosurfaceTechnologies 2016). For medical device-related applications, coupons made of polycarbonate, polyetheretherketone, stainless steel, polypropylene, glass or silicone are often used (Honraet, Goetghebeur and Nelis 2005, Nailis et al. 2010, Williams and Bloebaum 2010, Williams, Haymond, Beck, et al. 2012, Williams, Woodbury, Haymond, Parker and Bloebaum 2011). The majority of coupons that have been analyzed consist of relatively smooth, flat surfaces. In this study, a modified CDC biofilm reactor was developed such that it could hold coupons made of highly porous, bioabsorbable collagen (Figure 1B). The rationale for doing so was two-fold.

**Figure 1:**
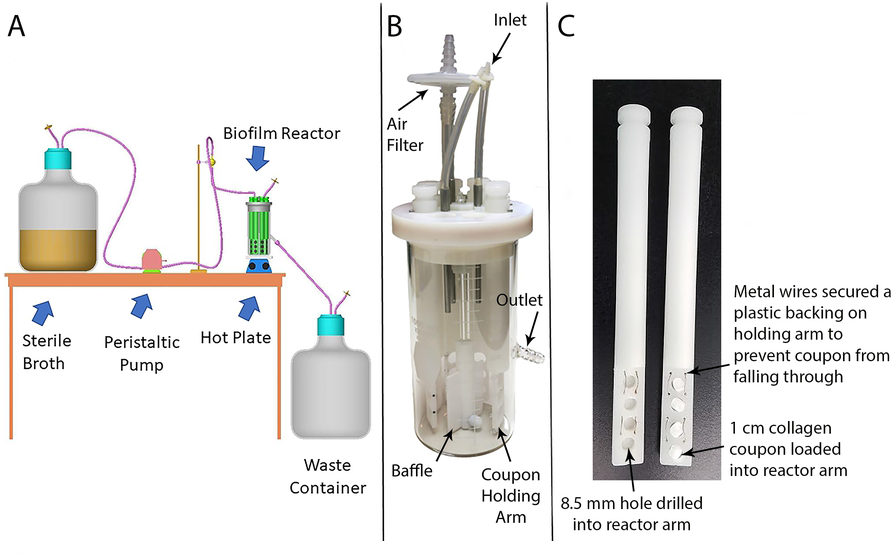
Design of the CDC biofilm reactor and modified holder. (A) Schematic of a general setup of a CDC biofilm reactor for biofilm growth. Source: Biosurface Technologies. (B) Components of a CDC biofilm reactor. (C) Customized holding arms into which collagen plugs were placed for biofilm growth. Labels detail modifications.

First, there is currently a paucity of data on antibiotic efficacy against biofilms that are grown on biologically relevant materials. Collagen constitutes 75% of the dry weight of human skin, and is a major component of extracellular matrix and multiple tissue types that have potential to be affected by biofilm formation (Shoulders and Raines 2009). It has also been shown that multiple bacterial species adhere to collagen, and may aid in their ability to colonize and potentially infect tissue types (Kang et al. 2013, Nallapareddy et al. 2000, Switalski et al. 1993, Vercellotti et al. 1985). The primary question to be answered in this study was whether biofilms that attached to biological material, such as highly porous and complex collagen, displayed similar antibiotic susceptibility profiles to biofilms that grew on relatively smooth, flat surfaces such as polycarbonate. As a secondary measure, the stability of bioabsorbable collagen in the modified reactor was assessed. More specifically, it needed to be confirmed that collagen would not disintegrate/absorb in the timeframe that biofilms were grown on it.

It was hypothesized that biofilms of varying species grown on a complex collagen network would be less susceptible to antibiotics as compared to biofilms grown on polycarbonate coupons. A secondary aspect of the study was that similar to a previously modified CDC biofilm reactor (Williams, Haymond, Beck, Savage, Chaudhary, Epperson, Kawaguchi and Bloebaum 2012, Williams, Haymond, et al. 2011, Williams, Haymond, Woodbury, et al. 2012, Williams et al. 2014, Williams, Woodbury, Haymond, Parker and Bloebaum 2011), the design of the holding rods in this system provide potential to broaden the scope of substrates on which biofilms can be grown in a CDC reactor for future *in vivo* analyses.

## Materials and Methods

### Reagents and Materials

Tryptic soy broth (TSB), brain heart infusion (BHI) broth, agar, cation-adjusted Mueller Hinton broth (CAMHB) and phosphate buffered saline (PBS) were purchased from Fisher Scientific (Waltham, MA). Blank polypropylene holders for the CDC reactor were purchased from Biosurface Technologies (Bozeman, MT). Collaform collagen plugs were purchased from Implant Logistics (La Crosse, WI). All antibiotics used were purchased from Sigma Aldrich (St. Louis, MI) or TCI America (Portland, OR). Antibiotics for each bacterium were chosen based on common clinical use. E-Test strips for MIC testing of amoxicillin and erythromycin were purchased from Biomérieux (Durham, NC).

### Bacterial Isolates

Bacterial isolates were chosen because of their use in various applications including standards assays and animal models, their known pathogenicity as well as their ability to form well-established biofilms (Luppens et al. 2002, Marshall et al. 1996, Peeters et al. 2008, Reimer et al. 1981). All isolates were purchased from the American Type Culture Collection (ATCC) and included *Staphylococcus aureus* ATCC 6538, *Pseudomonas aeruginosa* ATCC 27853, *Escherichia coli* ATCC 9637, *Acinetobacter baumannii* ATCC BAA 1605, *Bacillus subtilis* ATCC 19659, methicillin-resistant *S. aureus* (MRSA) USA 300, MRSA USA 400, *Streptococcus mutans* ATCC 25175, *S. epidermidis* ATCC 35984, and carbapenem-resistant *Klebsiella pneumoniae* ATCC BAA-1705. All were subbed on TSA or, in the case of some experiments with *S. mutans*, BHI agar, and incubated overnight at 37º C. Frozen stocks were maintained in BHI broth with 30% glycerol. Isolates were subbed and incubated 24 – 72 hours prior to inoculation in a biofilm reactor.

### Reactor and Materials Design

Standard polypropylene CDC biofilm reactor holders were used to hold polycarbonate coupons (Figure 1A). Custom holders were made for the collagen plug coupons (Figure 1B). To do so, four holes of 8.5 mm diameter each were drilled in the lower portion of a blank polypropylene holder (Figure 1B). This diameter was smaller than the diameter of standard reactor coupons (12.7 mm). The size difference was to ensure a tight fit of the collagen in the holders.

To make the collagen plugs, medical grade Collaform collagen was purchased as 1 cm x 2 cm plugs. The collagen was aseptically removed from packaging and cut into coupons with a sterile blade. Coupon size was 1 cm diameter x 0.3 cm height. Coupons were aseptically loaded into modified reactor arms (Figure 1B) that had already been autoclaved. The collagen coupons remained securely in place when exposed to the shear forces in the reactor. Set up of each reactor was performed inside a sanitized biosafety cabinet.

### Biofilm Growth

From a fresh culture, a 0.5 McFarland standard of each bacteria was made, which equated to ∼7.5 × 10^7^ colony forming units (CFU)/mL. One mL of the 0.5 McFarland solution was inoculated into 500 mL of BHI in the CDC biofilm reactor. The reactor was placed on a hot plate set at 34° C and a baffle rotation of 130 rpm for 24 h. After the 24 h batch growth, a continuous flow of 10% BHI was flowed through the reactor at ∼6.9 mL/min for an additional 24 h for a total of 48 h of growth.

### Baseline Quantification

To obtain a baseline of biofilm/coupon, two holding arms were aseptically removed from a reactor and rinsed in 1x PBS. An n=6 coupons were individually placed in a test tube that contained 2 mL CAMHB, vortexed for 1 min, sonicated for 10 min, then vortexed a final time for approximately 10 sec. A 10-fold dilution series was used to plate bacteria in duplicate on tryptic soy agar (TSA). Agar plates were incubated overnight at 37° C. CFU were counted and used to calculate the CFU/coupon.

### Antibiotic Treatment

Prior to assessing the antibiotic susceptibility of biofilms, the minimum inhibitory concentration (MIC) of each antibiotic was determined against each bacterial strain (see Table 2). For all antibiotics, except amoxicillin and erythromycin against *Streptococcus mutans*, a modified protocol of the Clinical and Laboratory Standards Institute (CLSI) guideline M100 was used. In short, using a fresh overnight culture of bacteria, a 0.5 McFarland standard was made in PBS using a nephelometer. The stock solution was diluted in PBS 1:100 to achieve a concentration of ∼7.5 × 10^5^ CFU/mL. A 96-well plate was set up such that a final volume of 100 µL was present in each well. Column 1 served as the negative control of growth (antibiotic only without bacteria added) and Column 11 served as the positive control of growth (bacteria only, no antibiotic).

To accomplish this, 100 µL of CAMHB that contained antibiotic only (64 µg/mL) was pipetted into each well of column 1 to serve as the negative control. Into columns 2-11, 50 µL of CAMHB were added to each well. Subsequently, 50 µL of CAMHB that contained a concentration of 256µg/mL antibiotic were added to each well of column 2 using a multi-channel pipet. The solution was mixed, and then 50 µL were removed and added to wells of column 3. This 1:2 dilution process was continued until column 10 and resulted in a range of antibiotic testing from 64 µg/mL to 0.0625 µg/mL. Lastly, into each well of columns 2-11, 50 µL of the bacterial solution were added, with column 11 serving as the positive control. The 96-well plate was covered with adhesive film and incubated 24 h at 37° C. The concentration of antibiotic that inhibited pellet formation or turbidity at the 24 h time point was considered the MIC. Once the MIC was determined, biofilm analysis was performed.

To determine the MIC of amoxicillin and erythromycin against *S. mutans*, an E-Test was used. In short, *S. mutans* was cultured on BHI agar and incubated for 48 hours under 5% CO_2_. A 1.0 McFarland standard solution was made in PBS resulting in 2.8 × 10^8^ CFU/mL solution. This was used to make lawns of bacteria on BHI agar by stroking back and forth with an inoculated Q-tip in three directions. After drying, E-Test strips for each compound were laid (n=2 per plate, 2 plates per compound). The plates were incubated for 24 hours under 5% CO_2_ and then analyzed per manufacturer’s instructions.

Following a reactor run, n=5 polycarbonate and n=6 collagen coupons were aseptically removed from a reactor, rinsed in 1x PBS, and each coupon was individually placed in a test tube that contained 2 mL solution of antibiotic in CAMHB. Each of the antibiotics were tested initially at 50, 100, and 200 µg/mL concentrations. In some instances, data spread needed to be resolved so additional concentrations were tested down to 25 or up to 400, or 600 µg/mL. Coupons were incubated for 24 h after which time each was quantified as described above.

### Scanning Electron Microscopy Imaging

Scanning electron microscopy (SEM) imaging was used to directly observe biofilm morphology and confirm growth on both coupon types. To perform SEM imaging, biofilms were grown following the same protocol as above. Notably, coupons used for SEM imaging to assess morphology were not the same coupons used for the antibiotic testing. This was not possible given the need to fix and process biofilms, which inherently leads to cell death and would have skewed the antibiotics or baseline quantification data.

To process for SEM imaging, a holder was aseptically removed from a reactor, rinsed one time in 1x PBS, and each coupon was individually submerged in modified Karnovsky’s fixative (2.5% glutaraldehyde and 4% formaldehyde in 0.2 M PBS, pH 7.4) for ∼1 h. The fixed specimens were dehydrated in 100% ethanol for ∼1 h, and then air-dried for ∼30 min.

Coupons were placed on an SEM stage and secured using double-sided carbon tape, then coated using a Hummer 6.2 gold sputter coater (Anatech LTD). All coupons were imaged with a JEOL JSM-6610 SEM in secondary electron emission mode.

In addition to assessing biofilm morphology, because a combination of vortexing and sonication was used to remove/disrupt biofilms as part of the baseline and treatment studies, it was important to confirm that the vortex and sonication process did in fact remove the large majority of biofilm from the surface of the coupons. After a thorough literature review, removal of biofilms from collagen using vortex and sonication does not appear to have been confirmed previously. Thus, for each bacterium, biofilms were grown on both materials, samples were rinsed, vortexed, and sonicated as described, and were imaged by SEM (n=3) to determine if that procedure effectively removes biofilm for an accurate quantification.

Lastly, in order to compare surface morphologies of the collagen and polycarbonate coupon materials without bacteria on them, n=3 new and unused (negative control) coupons were soaked in 10% BHI for 48 hrs. Coupons were then aseptically removed and processed as described above for SEM imaging.

### Statistical Analysis

Outcome measures (i.e. bacterial counts) were analyzed using an independent sample *t* test for comparisons with alpha level set at 0.05 throughout.

## Results

### Biofilm Growth/SEM Imaging

SEM images showed that the collagen material alone was amorphous with a polymeric strand network that had deep crevices throughout (Figure 2A-C). The polycarbonate surface had residual machining marks and isolated regions of raised material with ridges and undulation (Figure 2D-F). Data showed that biofilms of all species formed on both material types, with some variation in numbers (detailed below). Representative images of *P. aeruginosa* ATCC 27853, MRSA USA 400 and *S. mutans* ATCC 25175 are provided to show biofilm formation and morphologies (Figures 3, 4 & 5, respectively).

**Figure 2:**
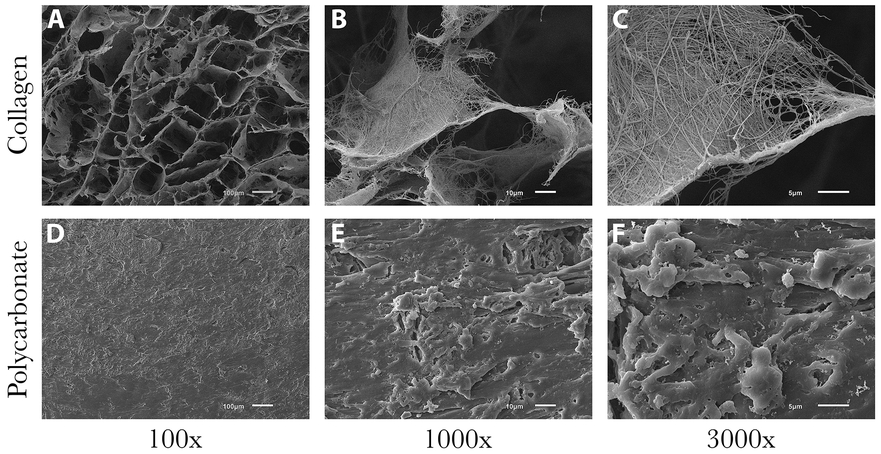
Representative SEM images of collagen and polycarbonate coupon surfaces. (A-C) Surface of a collagen coupon after soaking in broth only (no bacteria present). Deep valleys and ravines were consistent throughout the amorphous structure. (D-F) Surface of a polycarbonate coupon after soaking in broth only (no bacteria present). Ridges and plateaus had an undulating, yet mostly smooth surface, in particular relative to collagen.

SEM images showed that *P. aeruginosa* ATCC 27853 biofilms were able to form readily on both material types (Figure 3). This isolate did not form into plumes of three-dimensional structure. Rather, communities appeared to grow in stacked sheets (Figure 3). Higher-power images resolved extracellular matrix (ECM)-like fibers extending from one *P. aeruginosa* cell to another (Figure 3E & F).

**Figure 3:**
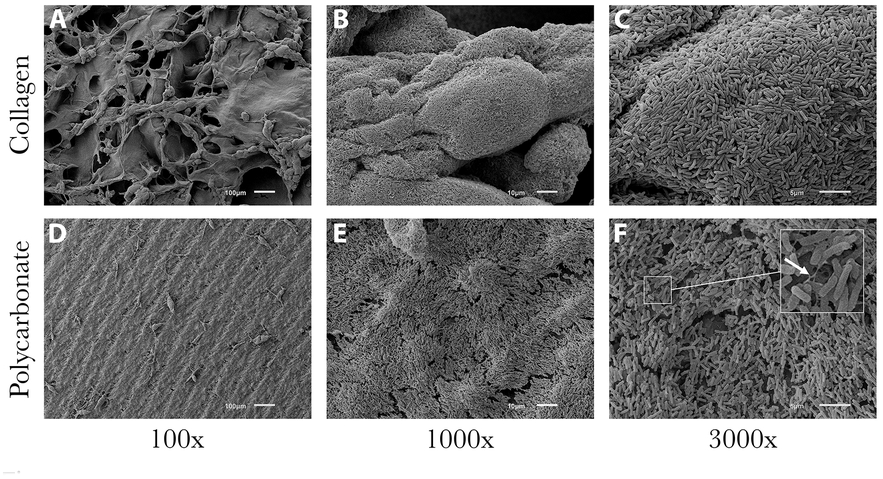
Representative SEM images of *P. aeruginosa* ATCC 27853 biofilm formation on both material types. (A-C) Surface of a collagen coupon that had biofilm coverage. Biofilm structure conformed to the polymeric collagen network. As noted in Figure 2, deep valleys and ravines were consistent throughout the amorphous structure. (D-F) Surface of a polycarbonate coupon on which biofilms of *P. aeruginosa* ATCC 27853 grew. Biofilm structure was estimated to be greater than 20 cell layers thick. Growth followed the contour of the surface, even displaying the coupon machine marks (100x). Arrow (panel F) indicates extracellular matrix components that were observed.

Staphylococcal biofilms produced three dimensional structures with vertical stacking morphology wherein cells grew on top of one another in sheet-like structures on collagen that delved deep into the collagen crevices (Figure 4A-C). On polycarbonate coupons, light plumes of biofilm formed primarily on the stiff, raised regions of the material surface (Figure 4D-F). On both material types, biofilm growth followed the contour of the material surface (see Figure 4). Extracellular matrix (ECM) development was sporadic, nevertheless fibers of matrix were seen throughout staphylococcal biofilms on both material types (Figure 2E & F). Biofilm growth was similar for all staphylococcal isolates (i.e., MRSA USA300, *S. aureus* and *S. epidermidis)*. *S. epidermidis* ATCC 35984 produced the most highly robust biofilms of any isolate tested, appearing extremely thick and canyon-like with visible water channels.

**Figure 4:**
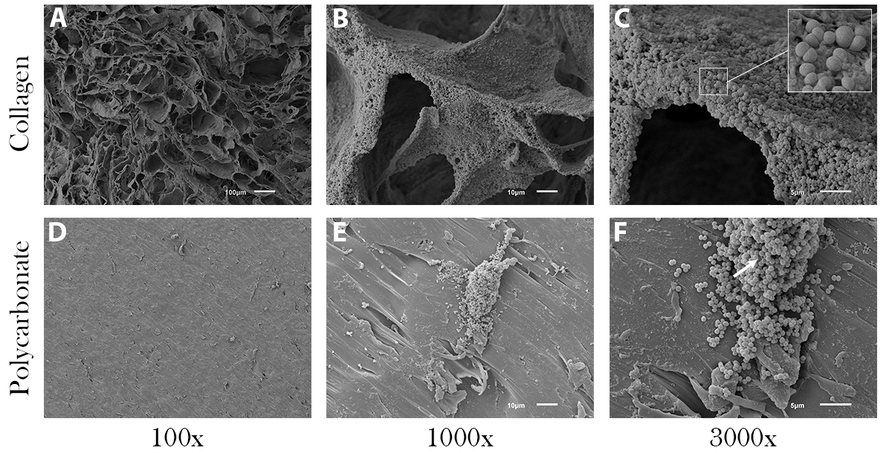
Representative SEM images of MRSA USA 400 biofilm on both material types. (A-C) Surface of collagen with biofilm growth. Biofilm structure resulted in uniform coverage, but did not appear to plume as other staphylococcal isolates did. Rather, MRSA USA 400 displayed sheet-like growth on collagen. (D-F) Surface of a polycarbonate coupon that had sparse coverage, although where biofilm did form, it plumed to an estimated 30 cell layers thick. Arrow (panel F) indicates a biofilm plume.

All other species produced biofilms with slight variations in thickness and coverage. In the case of *A. baumannii* ATCC BAA 1605, biofilms formed into sheet-like structures on both collagen and polycarbonate coupons. Isolated regions with matrix-like materials were observed. Layered sheet-like structures of biofilm were also seen in *E. coli* ATCC 6937 and *B. subtilis* ATCC 19659 with *B. subtilis* having the most significant ECM development on the smooth, flat polycarbonate coupons of all biofilms. Carbapenem-resistant *K. pneumoniae* ATCC BAA-1705 grew a dense, consistent smooth-layer biofilm with small amounts of ECM on polycarbonate. *S. mutans* ATCC 25175 grew a scattered monolayer biofilm on polycarbonate and thick plumes of biofilm covering most of the collagen (Figure 5). On both materials there was little ECM.

**Figure 5:**
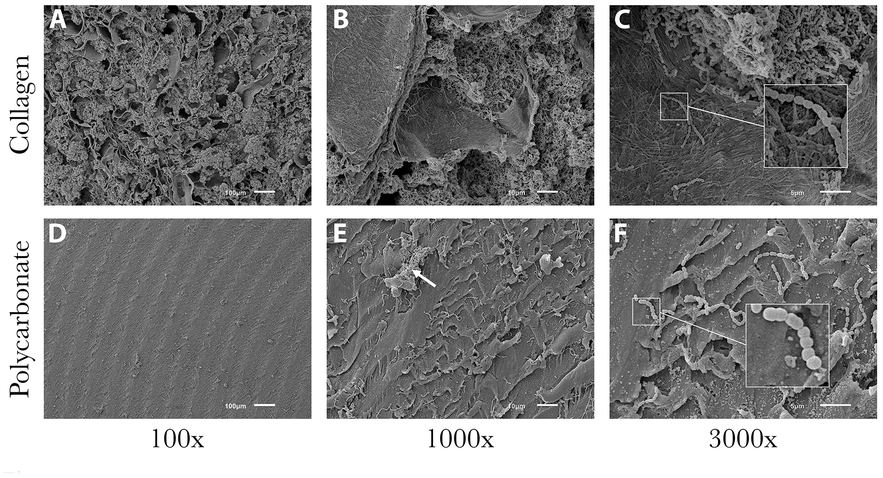
Representative SEM images of *S. mutans* ATCC 25175 biofilm formation on both material types. (A-C) Surface of collagen with biofilm coverage. Cells of *S. mutans* filled the ravines and crevices of the collagen material. (D-F) In contrast to collagen, *S. mutans* grew in small plumes on the surface of polycarbonate with sporadic coverage. Arrow (panel E) indicates a biofilm plume.

SEM images of coupons that had been vortexed and sonicated indicated that the large majority of biofilms had been removed from the surface (Figure 6). This confirmed that when quantifying the CFU/sample, the majority of bacterial cells were accounted for in the 10-fold dilution series. The degree of biofilm removal was similar for all biofilms (Figure 6), with the exception of *S. epidermidis* ATCC 35984 (Figure 7). Although there was still a relatively fair amount of *S. epidermidis* ATCC 35984 cells that remained on each material type post-vortex/sonication, the levels of biofilm that were present initially were far more than any other isolate (see Figures 3-7). Thus, the reduction was still significant. Based on surface area coverage and reduction of the biofilm layers to a monolayer of cells, it was estimated that less than 5% of cells remained on the surface after vortexing and sonication.

**Figure 6:**
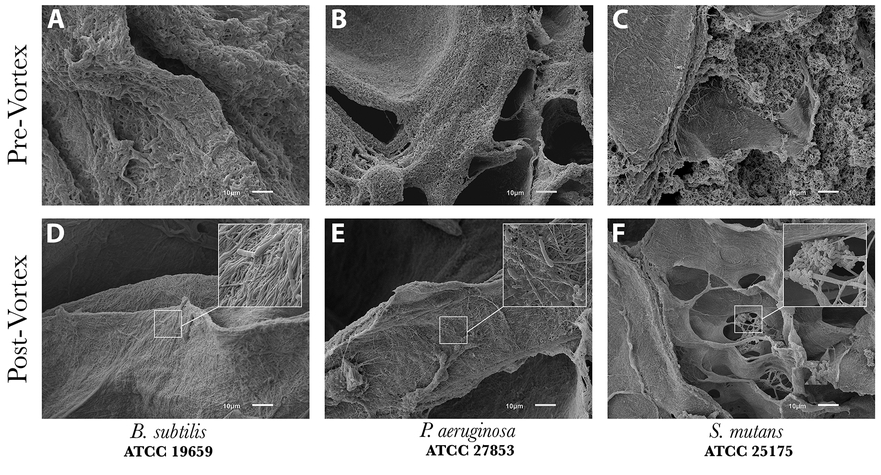
Representative SEM images of collagen coupons with three isolates pre-and post-vortex/sonication to demonstrate the ability of the process to remove bioburden from material surfaces. (A-C) Representative images of biofilm structures prior to vortex and sonication. (D-F) Images of residual cells on collagen after vortex and sonication. Data indicated that biofilms were effectively disrupted/removed with >5% of cells remaining on the surface.

**Figure 7:**
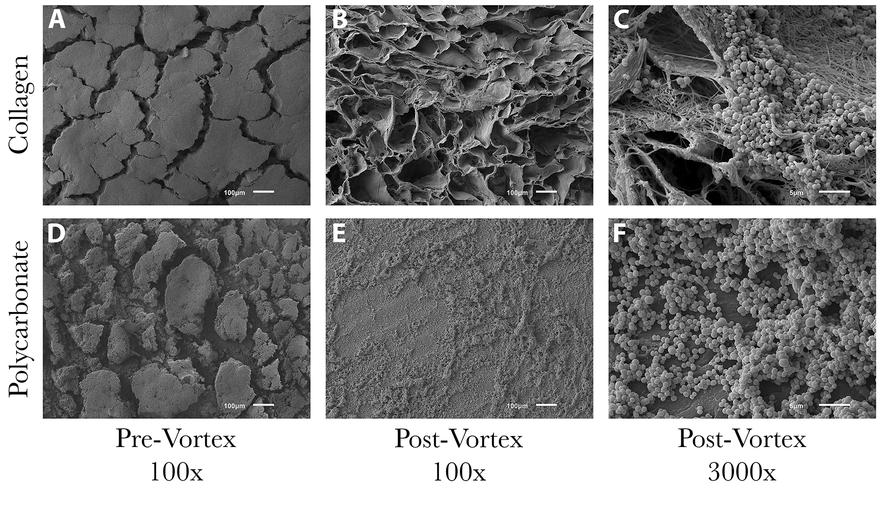
Representative SEM images of *S. epidermidis* ATCC 35984 biofilms on collagen and polycarbonate coupons pre-and post-vortex/sonication. (A) Heavy amounts of biofilm formed on collagen coupons, making the surface morphology unobservable. (B-C) Images showed that the large majority of biofilm burden had been removed by vortex/sonication, but clusters of cells still remained. (D) Similar to collagen, large plumes of biofilms formed on polycarbonate coupons with deep ravines. Although these ravines may have formed during the dehydration procedure, it is likely they were sites of water channels that provided fracture points within the biofilm communities. (E-F) Images showed that there was still a fair amount of surface coverage by biofilms post-vortex/sonication. However, it was estimated that there were >5% of cells remaining on the surface similar to other bacterial isolates examined.

### Baseline Quantification and Antibiotic Treatment

Quantification data of untreated coupons (positive controls of growth) is reported in Table 1. Of the ten isolates examined, six had growth profiles on collagen versus polycarbonate that were statistically significantly different (Table 1). In each case where there was a difference, more growth was present on collagen compared to polycarbonate (Table 1).

**Table 1:**
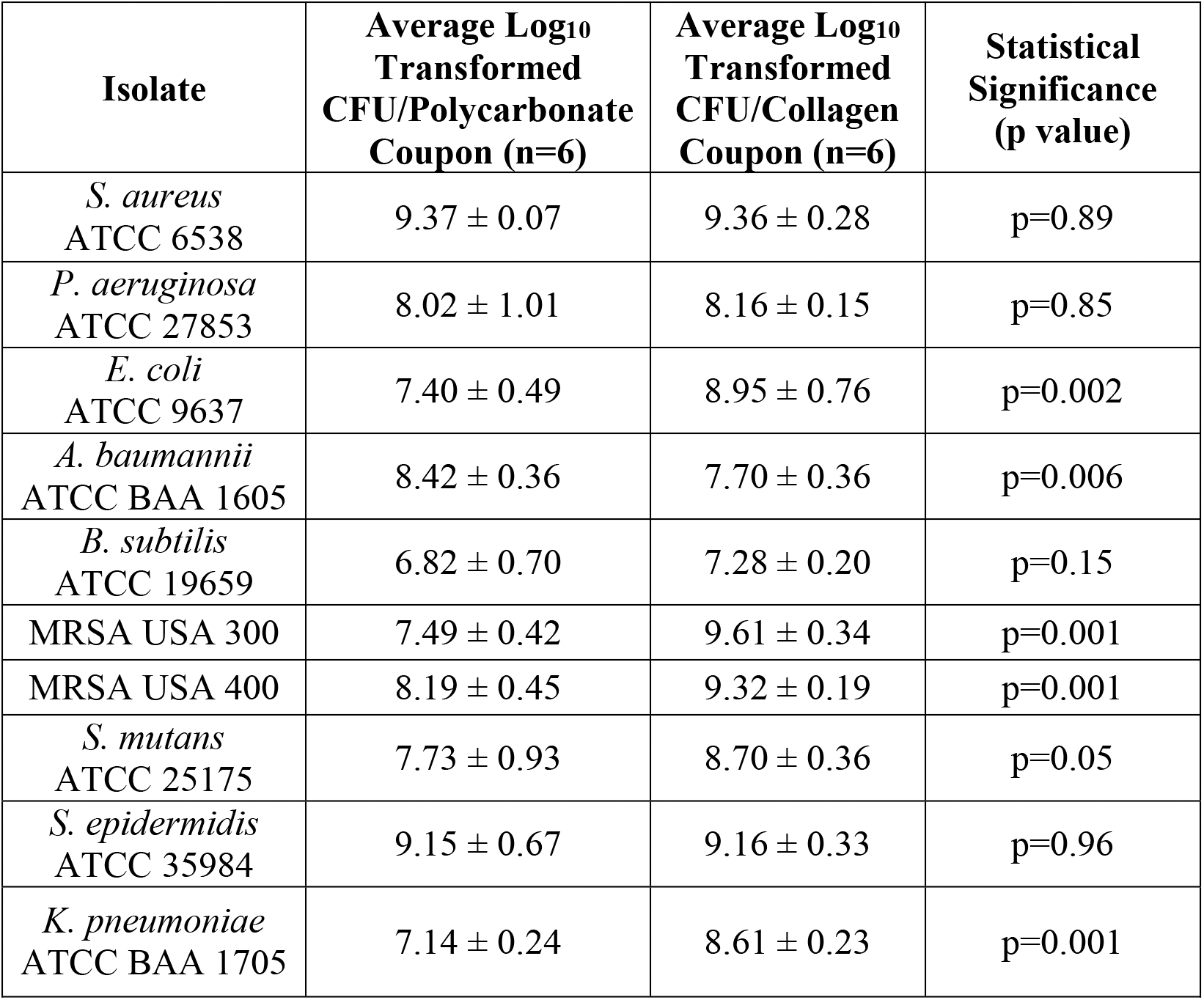
Baseline (control) quantifications of coupons.

| **Isolate** | **Average Log<sub>10</sub>**<br>**Transformed**<br>**CFU/Polycarbonate**<br>**Coupon (n=6)** | **Average Log<sub>10</sub>**<br>**Transformed**<br>**CFU/Collagen**<br>**Coupon (n=6)** | **Statistical**<br>**Significance**<br>**(p value)** |
| --- | --- | --- | --- |
| *S. aureus*<br>ATCC 6538 | 9.37 ± 0.07 | 9.36 ± 0.28 | p=0.89 |
| *P. aeruginosa*<br>ATCC 27853 | 8.02 ± 1.01 | 8.16 ± 0.15 | p=0.85 |
| *E. coli*<br>ATCC 9637 | 7.40 ± 0.49 | 8.95 ± 0.76 | p=0.002 |
| *A. baumannii*<br>ATCC BAA 1605 | 8.42 ± 0.36 | 7.70 ± 0.36 | p=0.006 |
| *B. subtilis*<br>ATCC 19659 | 6.82 ± 0.70 | 7.28 ± 0.20 | p=0.15 |
| MRSA USA 300 | 7.49 ± 0.42 | 9.61 ± 0.34 | p=0.001 |
| MRSA USA 400 | 8.19 ± 0.45 | 9.32 ± 0.19 | p=0.001 |
| *S. mutans*<br>ATCC 25175 | 7.73 ± 0.93 | 8.70 ± 0.36 | p=0.05 |
| *S. epidermidis*<br>ATCC 35984 | 9.15 ± 0.67 | 9.16 ± 0.33 | p=0.96 |
| *K. pneumoniae*<br>ATCC BAA 1705 | 7.14 ± 0.24 | 8.61 ± 0.23 | p=0.001 |

MICs for each antibiotic against the bacterial species are reported in Table 2. Only one isolate (*K. pneumoniae*) was resistant to the antibiotics tested. Despite lack of susceptibility against *K. pneumoniae*, data are still provided to support the test method, i.e., to show that if an isolate was not susceptible, no biofilm reduction would have be present, but when susceptible, reduction in biofilm numbers could be considered accurate. Biofilm susceptibility outcomes for both material types are summarized below. Full results are provided in Table 3.

**Table 2:** Bacteria and the antibiotics they were tested against, including MIC for each.

| Bacteria | Antibiotic | MIC (µg/mL) |
| --- | --- | --- |
| *Staphylococcus aureus* ATCC 6538 | Ciprofloxacin | 0.25 |
|  | Cefazolin | ≤0.0625 |
| *Pseudomonas aeruginosa* ATCC 27853 | Tobramycin | 0.25 |
|  | Ciprofloxacin | 0.5 |
| *Escherichia coli* ATCC 9637 | Ceftriaxone | 0.0625 |
|  | Ciprofloxacin | ≤0.0625 |
| *Acinetobacter baumannii* ATCC BAA 1605 | Imipenem | 16 |
|  | Colistin | 1 |
| *Bacillus subtilis* ATCC 19659 | Ciprofloxacin | 0.5 |
|  | Vancomycin | 0.25 |
| Methicillin-resistant *Staphylococcus aureus* (MRSA) USA 300 | Vancomycin | 2 |
|  | Daptomycin | 8 |
| Methicillin-resistant *Staphylococcus aureus* (MRSA) USA 400 | Vancomycin | 1 |
|  | Daptomycin | 8 |
| *Streptococcus mutans* ATCC 25175 | Erythromycin | 0.047 |
|  | Amoxicillin | 0.064 |
| *Staphylococcus epidermidis* ATCC 35984 | Vancomycin | 1 |
|  | Ciprofloxacin | 0.5 |
| Carbapenem-resistant *Klebsiella pneumoniae* ATCC BAA-1705 | Ceftriaxone | >250 |
|  | Imipenem/Cilastatin | 32 |

**Table 3:** Average of Log_10_ transformed CFU/sample for each species and material at each concentration of antibiotic tested. Unless otherwise noted (at times coupons fell out of the holding arm in the reactor and were not included in analysis), n=6 collagen coupons were analyzed and n=5 polycarbonate coupons were analyzed.

| Bacteria | Antibiotic | Material | 25 µg/mL | 50 µg/mL | 100 µg/mL | 200 µg/mL | 400 µg/mL | 600 µg/mL | Hypothesis supported? (Y/N) |
| --- | --- | --- | --- | --- | --- | --- | --- | --- | --- |
| *S. aureus*<br>ATCC 6538 | Ciprofloxacin | Collagen | - | 8.89 ± 0.15 | 8.62±0.07 | 8.50±0.13 | 8.35±0.16 | - | Yes |
|  |  | Polycarbonate | - | 8.46±0.45 | 8.50±0.21 | 7.47±0.29 | 8.31±0.15 | - |  |
|  | Cephazolin | Collagen | - | - | 9.34±0.27 | 8.90±0.21 | 9.02±0.12 | - | No |
|  |  | Polycarbonate | - | - | 9.13±0.83 | 8.90±0.17 | 8.72±0.08 | - |  |
| *P. aeruginosa*<br>ATCC 27853 | Tobramycin | Collagen | 5.26±0.89 | *3.99±0.33 | §0±0 | - | - | - | Yes |
|  |  | Polycarbonate | 4.88±0.33 | 1.72±1.80 | *0±0 | - | - | - |  |
|  | Ciprofloxacin | Collagen | - | 4.29±0.76 | 4.64±0.90 | 4.12±0.49 | - | - | Yes |
|  |  | Polycarbonate | - | 4.11±0.34 | 3.87±0.74 | 1.77±1.62 | - | - |  |
| *E. coli*<br>ATCC 9637 | Ceftriaxone | Collagen | - | §6.32±1.21 | §3.60±3.65 | 4.17±3.53 | - | - | Yes |
|  |  | Polycarbonate | - | 0±0 | *0±0 | 0±0 | - | - |  |
|  | Ciprofloxacin | Collagen | - | 6.15±0.84 | §7.13±0.33 | §6.63±0.62 | - | - | Yes |
|  |  | Polycarbonate | - | 1.83±2.53 | 5.35±0.41 | 4.72±0.53 | - | - |  |
| *A. baumannii*<br>ATCC BAA 1605 | Imipenem | Collagen | - | 8.38±0.12 | 7.31±0.40 | 7.31±0.38 | - | - | No |
|  |  | Polycarbonate | - | 8.42±0.05 | 7.54±0.16 | 7.20±0.24 | - | - |  |
|  | Colistin | Collagen | - | 4.91±2.57 | 0±0 | 0±0 | - | - | No |
|  |  | Polycarbonate | - | 4.28±2.61 | 0±0 | 0±0 | - | - |  |
| *B. subtilis*<br>ATCC 19659 | Ciprofloxacin | Collagen | - | 5.71±0.64 | 3.99±0.35 | 0±0 | - | - | Yes |
|  |  | Polycarbonate | - | 5.58±0.76 | 0±0 | 0±0 | - | - |  |
|  | Vancomycin | Collagen | - | 5.68±0.46 | 4.64±0.36 | 5.13±0.41 | - | - | Yes |
|  |  | Polycarbonate | - | 5.57±0.31 | 0±0 | 1.84±2.52 | - | - |  |
| MRSA<br>USA 300 | Vancomycin | Collagen | - | - | 8.74±0.49 | 9.42±0.46 | 9.27±0.07 | - | Yes |
|  |  | Polycarbonate | - | - | 6.61±0.48 | 6.52±0.68 | 2.42±2.24 | - |  |
|  | Daptomycin | Collagen | - | 7.41±0.70 | 7.12±0.35 | 1.30±1.43 | - | - | Yes |
|  |  | Polycarbonate | - | 2.22±2.14 | 1.93±1.79 | 0±0 | - | - |  |
| MRSA<br>USA 400 | Daptomycin | Collagen | - | 7.48±0.25 | 6.68±0.53 | 5.50±0.83 | - | - | Yes |
|  |  | Polycarbonate | - | 4.48±1.03 | *3.27±1.03 | 1.62±2.36 | - | - |  |
|  | Vancomycin | Collagen | - | 8.87±0.37 | 9.00±0.19 | 8.78±0.50 | - | - | Yes |
|  |  | Polycarbonate | - | 6.54±0.42 | 6.48±0.56 | 7.17±1.05 | - | - |  |
| *S. mutans*<br>ATCC 25175 | Erythromycin | Collagen | - | 7.56±0.26 | 8.50±0.53 | 8.50±0.67 | - | - | No |
|  |  | Polycarbonate | - | 5.61±0.15 | 8.97±0.18 | 9.08±0.69 | - | - |  |
|  | Amoxicillin | Collagen | - | 8.05±0.20 | 8.13±0.39 | 7.77±0.23 | - | - | Yes |
|  |  | Polycarbonate | - | 4.98±0.33 | 4.81±0.38 | 5.23±0.28 | - | - |  |
| *S. epidermidis*<br>ATCC 35984 | Vancomycin | Collagen | - | 8.33±0.14 | §8.45±0.16 | 8.47±0.09 | - | - | No |
|  |  | Polycarbonate | - | 8.66±0.35 | 8.62±0.10 | 8.49±0.15 | - | - |  |
|  | Ciprofloxacin | Collagen | - | 8.32±0.17 | 7.46±0.06 | 6.34±0.05 | - | - | Yes |
|  |  | Polycarbonate | - | 8.55±0.10 | 6.76±0.34 | 6.07±0.11 | - | - |  |
| *K. pneumoniae*<br>ATCC BAA-1705 | Ceftriaxone | Collagen | - | 9.58±0.07 | 8.84±0.51 | 9.23±0.14 | 8.99±0.15 | 8.74±0.34 | N/A |
|  |  | Polycarbonate | - | 9.19±0.15 | 8.90±0.42 | 8.91±0.27 | 8.37±0.18 | *8.08±0.16 |  |
|  | Imipenem/<br>Cilastatin | Collagen | - | 9.29±0.07 | 9.63±0.57 | 9.23±0.18 | 9.10±0.09 | 8.97±0.10 | N/A |
|  |  | Polycarbonate | - | 8.74±0.15 | 9.41±0.25 | 8.84±0.23 | 8.82±0.16 | 8.75±0.21 |  |

\* n=4 coupons tested
§ n=5 coupons tested

Biofilms of *S. aureus* ATCC 6538 grown on collagen or polycarbonate were minimally affected by ciprofloxacin, but did show a statistically significant difference with biofilms on collagen having lower reduction than those on polycarbonate at 200 µg/mL (p=0.001; see Table 3). Susceptibilities were the same when exposed to cefazolin across a range of concentrations (Table 3; p>0.3 in both cases). In this data set, the hypothesis was supported in one instance with ciprofloxacin, but in no cases with cefazolin.

In the case of *P. aeruginosa* ATCC 27853, there was an approximately 4-5 Log_10_ reduction following exposure to ciprofloxacin at 50, 100, and 200 µg/mL (Table 3). Susceptibility of biofilms to ciprofloxacin on collagen was significantly less compared to those on polycarbonate (e.g., p=0.03 at 200 µg/mL). Biofilm susceptibility to tobramycin had similar outcomes. Tobramycin resulted in a 4-5 Log_10_ reduction at 50 µg/mL and complete kill at 100 µg/mL for both coupon types, with susceptibility being greater on polycarbonate coupons compared to collagen (Table 3; e.g., p=0.046 at 50 µg/mL). The hypothesis was supported with both antibiotics in this data set.

Biofilms of *E. coli* ATCC 9637 had the most notable differences in susceptibility profiles between collagen and polycarbonate growth (Table 3). In all data sets, both ciprofloxacin and ceftriaxone were more effective against biofilms on polycarbonate than those on collagen with ceftriaxone having more polarized effect than ciprofloxacin (Table 3). Differences between collagen and polycarbonate testing were all statistically significantly different. As a representative example, p=0.001 for ciprofloxacin at 100 µg/mL. The hypothesis was most strongly supported in this data set, in particular with ceftriaxone.

Biofilms of *A. baumannii* ATCC BAA 1605 were highly susceptible to colistin, with an 8 Log_10_ reduction (undetectable growth) on both collagen and polycarbonate coupons at 100 µg/mL, and near complete kill at 200 µg/mL (Table 3). The difference in CFU/coupon was significantly different from baseline controls (p<0.005 in all cases), but there was no statistically significant difference in the number of CFU/coupon between coupon types treated with colistin at 50 µg/mL (p=0.371). These results indicated that efficacy of colistin against *A. baumannii* biofilms was similar on both material types. Imipenem had minimal effect on biofilms on either material type up to 200 µg/mL, and indicated that biofilm susceptibility was similar on both materials (p=0.590). The hypothesis was not supported in any case for *A. baumannii* and the antibiotics tested.

Biofilms of *B. subtilis* ATCC 19659 were found to be more susceptible to vancomycin on polycarbonate than collagen (Table 3; e.g., p=0.001 at 100 µg/mL). Although there was no detectable growth of biofilms on polycarbonate exposed to vancomycin at 100 µg/mL, an anomaly was observed on polycarbonate coupons treated with vancomycin at 200 µg/mL; two of five coupons had growth. The experiment was repeated with similar outcomes, resulting in a large standard deviation at that concentration (Table 3). Biofilms on polycarbonate treated with ciprofloxacin were reduced by >6.5 Log_10_ units (undetectable growth) at 100 µg/mL, whereas those on collagen were reduced by only ∼3 Log_10_ units (p=0.001). At 200 µg/mL, ciprofloxacin eradicated biofilms completely on both polycarbonate and collagen. At 50 µg/mL, vancomycin and ciprofloxacin each reduced biofilms by ∼1 Log_10_ unit on polycarbonate and collagen coupons (p>0.1 in all cases; Table 3). At concentrations above 50 µg/mL, the hypothesis was supported for both antibiotics tested in this data set.

In the case of MRSA USA 300, daptomycin was more effective at eradicating biofilms on polycarbonate compared to collagen (p<0.001 in all cases), in particular at 400 µg/mL (Table 3). Vancomycin was nearly ineffective against biofilms on collagen (see Table 3) and reduced biofilms on polycarbonate coupons to a much greater degree (e.g., p=0.002 at 400 µg/mL).

Similar to USA300 data, daptomycin was more effective at eradicating MRSA USA 400 biofilms on polycarbonate compared to collagen (e.g., p=0.003 at 100 µg/mL). Biofilms on collagen showed no significant reduction by vancomycin at any concentration (Table 3) and on polycarbonate showed a reduction of approximately 1.5 Log_10_ units by vancomycin at 100 µg/mL, which was significant (p=0.001). Outcomes indicated that vancomycin was more effective at eradicating MRSA USA 400 biofilms on polycarbonate compared to collagen.

Biofilms of *S. mutans* ATCC 25175 on collagen and polycarbonate showed no significant reduction from baseline growth and no difference when exposed to 200 µg/mL erythromycin (p=0.198), indicating that erythromycin was equally ineffective at eradicating *S. mutans* biofilm on both material types. Biofilms on collagen were minimally affected by amoxicillin (Table 3.) However, biofilms on polycarbonate showed significant reductions of approximately 2.5 Log_10_ units by amoxicillin at all concentrations (e.g., p=0.001 at 200 µg/mL). These data indicated that amoxicillin was more effective at eradicating *S. mutans* biofilm on polycarbonate compared to collagen, which contrasted the results of erythromycin. Thus, outcomes with amoxicillin supported the hypothesis, whereas erythromycin did not.

Biofilms of *S. epidermidis* ATCC 35984 showed minimal reduction against biofilms on collagen or polycarbonate when treated with vancomycin, indicating that vancomycin was equally ineffective at eradicating *S. epidermidis* biofilms on both material types. Biofilms on collagen showed a reduction of approximately 1.5 Log_10_ units by ciprofloxacin at 100 µg/mL and approximately 2.5 Log_10_ units at 200 µg/mL (Table 3). Biofilms on polycarbonate were reduced by approximately 2.5 Log_10_ units with ciprofloxacin at 100 µg/mL and approximately 3 Log_10_ units at 200 µg/mL. There was a statistical difference between collagen and polycarbonate outcomes with ciprofloxacin (e.g., p=0.004 at 200 µg/mL). Thus, data indicated that ciprofloxacin was more effective against *S. epidermidis* biofilms on polycarbonate compared to those on collagen. Outcomes supported the hypothesis with ciprofloxacin, but not vancomycin.

In the case of *K. pneumoniae* ATCC BAA-1705, ceftriaxone and imipenem/cilastatin had minimal effect against biofilms on collagen or polycarbonate. There was about 1 Log_10_ more bacteria that grew on collagen compared to polycarbonate. In all cases, there were more bacteria on coupons in treatment groups than there were on positive controls. These data neither supported nor unsupported the hypothesis as the isolate was resistant to both antibiotics, yet were included to provide a comparison of outcomes when susceptibility was not present.

Taken together, data indicated that in 13/18 cases the hypothesis was supported. *K. pneumoniae* data were not included in the outcome as biofilms treated with antibiotics had more growth than positive controls.

## Discussion

Biofilms can be grown on a wide variety of surfaces, materials and exposed to myriad environmental conditions. One of the most common reactor systems to grow biofilms is the CDC biofilm reactor, which is beneficial in that it is a robust system that can be modified to hold a broad variety of coupon types (Buckingham-Meyer, Goeres and Hamilton 2007, Goeres, Loetterle, Hamilton, Murga, Kirby and Donlan 2005, Williams, Woodbury, Haymond, Parker and Bloebaum 2011). As such, its utility spans a broad scope of research and development. In this study, the antibiotic susceptibility of multiple bacterial species was assessed with specific focus on determining whether biofilm growth on coupons made of a complex collagen material or on a relatively smooth, non-complex polycarbonate surface would influence susceptibility. The rationale was, there is significant literature on susceptibility profiles of biofilms grown on hard, relatively smooth polymer surfaces, but a paucity of data for susceptibility on more complex, biologically-relevant materials such as collagen. Yet it is important to understand susceptibility profiles given that clinical antibiotics used in human applications may find utility in targeting biofilms that have significant and direct contact with soft and hard tissue, a primary constituent of which is collagen (Fratzl et al. 2004, Lovell et al. 1987, Neuman and Logan 1950).

One component of the study was to determine whether bioabsorbable collagen would dissolve/disintegrate in the modified CDC biofilm reactor. Results indicated that the collagen did not dissolve/disintegrate within the time frame of growth. Indeed, additional data further suggested that collagen was stable to culture conditions for as long as 8 days in the reactor. This demonstrated that growth on bioabsorbable collagen could be performed in the CDC reactor and allows for future applications of this collagen biofilm system toward *in vitro* and *in vivo* applications. For example, in a diabetic pig wound study, biofilms are currently being grown on collagen coupons for inoculation and analysis.

Biofilm susceptibility data indicated that in the majority of instances (13/18), the general hypothesis—that biofilms grown on polycarbonate would be more susceptible to antibiotics than those grown on collagen—was supported. Pairings that supported the hypothesis were ciprofloxacin at 200 µg/mL against *S. aureus* ATCC 6538, tobramycin and ciprofloxacin against *P. aeruginosa* ATCC 27853, ceftriaxone and ciprofloxacin against *E. coli* ATCC 9637, vancomycin and ciprofloxacin against *B. subtilis* ATCC 19659, vancomycin and daptomycin against the methicillin-resistant *S. aureus* strains USA 300 and USA 400, amoxicillin against *S. mutans* ATCC 25175, and ciprofloxacin at 100 µg/mL against *S. epidermidis* ATCC 35984. Pairings that did not support the hypothesis were cefazolin against *S. aureus* ATCC 6538, colistin and imipenem against *A. baumannii* ATCC BAA 1605, erythromycin against *S. mutans* ATCC 25175, and vancomycin against *S. epidermidis* ATCC 35984. Carbapenem-resistant *K. pneumoniae* ATCC BAA 1705 was resistant to both ceftriaxone and imipenem/cilastatin.

Baseline quantification data indicated that for five of the ten isolates, both coupon types had similar amounts of biofilm. The five that did not were, *E. coli*, MRSA USA300, MRSA USA400, *S. mutans*, and *K. pneumoniae*, which formed biofilms more readily and to a greater degree on collagen coupons compared to polycarbonate. In four of those cases (*E. coli*, MRSA 300, MRSA 400, and *S. mutans*), reduction of biofilms on polycarbonate coupons was significantly greater than on collagen coupons by one or both antibiotics. These data highlighted the importance of confirming efficacy profiles of antibiotics against biofilms on material types that may be relevant to a given application. As in the majority of instances biofilms were more tolerant to antibiotics on collagen than on hard surface polycarbonate, for applications of clinical relevance, it may be beneficial to assess antibiotic biofilm efficacy profiles against an isolate(s) in the presence of biologically relevant materials.

Limitations of the study included a single time of biofilm growth (48 hr). This was consistent with published standards. However, future work could include the addition of growing biofilms for a longer period of time to increase maturity. Biofilms that develop over a longer period of time may more closely replicate biofilms in clinical environments and thus potentially influence susceptibility profiles (Wolcott et al. 2010). Time and concentration of antibiotic exposure can also be analyzed in future work to determine what influence these have on more significant eradication of biofilms.

Taken together, data from eight isolates (thirteen isolate/antibiotic pairings) supported the hypothesis, that susceptibility to antibiotic would be greater on polycarbonate coupons compared to collagen. In the case of *A. baumannii* ATCC BAA 1605, antibiotics were able to eradicate biofilms to the same degree whether grown on collagen or polycarbonate coupons. *K. pneumoniae* ATCC BAA 1705 was excluded from susceptibility analysis as it was resistant to the antibiotics tested.

Higher concentrations of antibiotic were not tested given that the primary outcome measure was to determine if susceptibility was similar on two material types, not to determine the concentration of antibiotic that would reduce biofilms to a specific level. Although the reduction of biofilms was not a primary outcome measure, if data were to be applied to clinically relevant paradigms, this would be an important and application-dependent factor.

In conclusion, susceptibility of biofilms to antibiotics was significantly different in the majority of instances following 48 h of growth on biologically relevant collagen or polycarbonate material. Although there are only speculative reasons why some antibiotic-bacterial pairing produce significant differences in biofilm susceptibility on different substrates, this study demonstrates that there can be differences in susceptibility. On a go-forward basis, these data are valuable for advancing information on antibiotic susceptibility testing of biofilms grown on a variety of material types and encourage the independent evaluation of susceptibilities on study-relevant materials.

## Acknowledgments

This material is the result of work supported with resources and the use of facilities at the George E. Wahlen Department of Veterans Affairs. The contents do not represent the views of the U.S. Department of Veterans Affairs or the United States Government. The authors thank Seungah Goo, Li Cadenas, Marissa Badham, Scott Porter, Brooke Kawaguchi and Nick Taylor for their technical support.

